# Illumina sequencing analysis of the ruminal microbiota in high-yield and low-yield lactating dairy cows

**DOI:** 10.1101/325118

**Authors:** Jinjin Tong, Hua Zhang, Delian Yang, Benhai Xiong, Linshu Jiang

## Abstract

In this study, differences in the ruminal bacterial community between high-yield and low-yield lactating dairy cows fed the same diets were investigated. Sixteen lactating dairy cows with similar parity were divided into two groups based on their milk yield: high-yield (HY) and low-yield (LY) groups. On day 21, rumen content samples were collected, and the microbiota composition was determined using Illumina MiSeq sequencing of the 16S rRNA gene. During the study period, dry matter intake (DMI) and milk yield were measured daily, and milk composition was assessed 3 times per week. The results showed that the milk of the LY group tended to have higher fat (*P*=0.08), protein (*P*=0.01) and total solid (*P*=0.04) contents than that of the HY group, though the HY group had higher ruminal acetate (*P*=0.05), propionate (*P*=0.02) and volatile fatty acid (VFA) (*P*=0.02) concentrations. Principal coordinate analysis indicated significant differences in ruminal bacterial community composition and structure between the HY group and LY group. Overall, Bacteroidetes (HY group: 52.91±3.06%; LY group: 61.88±3.03%) was the predominant phylum, followed by Firmicutes (HY group: 41.10±2.74%; LY group: 32.11±2.97%). The abundances of *Ruminococcus 2*, Lachnospiraceae and *Eubacterium coprostanoligenes* were significantly higher in the HY group than in the LY group. In addition, 3 genera—*Anaerostipes*, *Bacteroidales* and *Anaeroplasma*—were identified as biomarker species with the greatest impacts on the ruminal community structure in the LY group. These findings facilitate the understanding of bacterial synthesis within the rumen and reveal an important mechanism underlying differences in milk production in dairy cows.

## Introduction

A symbiotic relationship exists with regard to the rumen microbiota of cattle. The rumen is a highly specialised organ of ruminant animals that promotes a community of mutualistic microbial species while simultaneously absorbing nutrients derived from digestion of plant fibre and cellular material [1]. In dairy cows, the rumen microbial community plays a critical role in the volatile fatty acid (VFA) production, B-vitamin synthesis, and microbial protein biosynthesis, which are critical for the animal’s well-being and efficient milk production [2]. Moreover, the structure of the bacterial community has been correlated with the production traits [3], production variables [4–7], and milk production and composition [8] of dairy cows. It has been reported that rumen microbial dynamics involve both core and variable microbiotal components [9–11]. In addition, similar to the microbial community in the gut of non-ruminants, the structure and function of the microbial community in the cow rumen are shaped by dynamic physical, chemical, and predatory environments. In turn, the microbial community regulates nutrient cycling to the host [12]. However, a more in depth comparison is warranted to improve our understanding of differences in rumen bacterial community composition between high-yield and low-yield dairy cows.

Recent efforts to study the rumen microbiome have focused on identifying and quantifying ruminal microbial communities [8, 13]. As a powerful molecular approach for taxonomic analyses, the application of 16S rRNA gene sequencing technology has provided novel insight into the microbiome ecology of gastrointestinal tracts [14, 15]. Indeed, this technique has been widely used to study microbial diversity and the metabolic capabilities of microbiomes in different ecological niches [16], fermented food [17, 18], waste-water treatment facilities [19], and human and animal gastrointestinal tracts [20–22]. The objective of the present study was to examine differences in ruminal bacterial community compositions between high-yield and low-yield lactating cows.

## Materials and Methods

### Animals and Experimental Design

The experimental protocol was approved by the Institutional Animal Care and Use Committee at the Beijing University of Agriculture, in compliance with regulations for the administration of affairs concerning experimental animals (The State Science and Technology Commission of P. R. China, 1988). According to the principle of parity and lactation days, 16 Holstein lactating dairy cows of similar parity were used and assigned to a high-yield group (average production 31.90±1.76 kg/d, mean±SD) or a low-yield group (average production 19.30±1.76 kg/d), with 8 each. The test period was 21 d, with a pre-feeding period of 14 d and a treatment period of 7 d. These lactating dairy cows were the fed same diet, the composition of which is shown in Table 1.

**Table 1.**
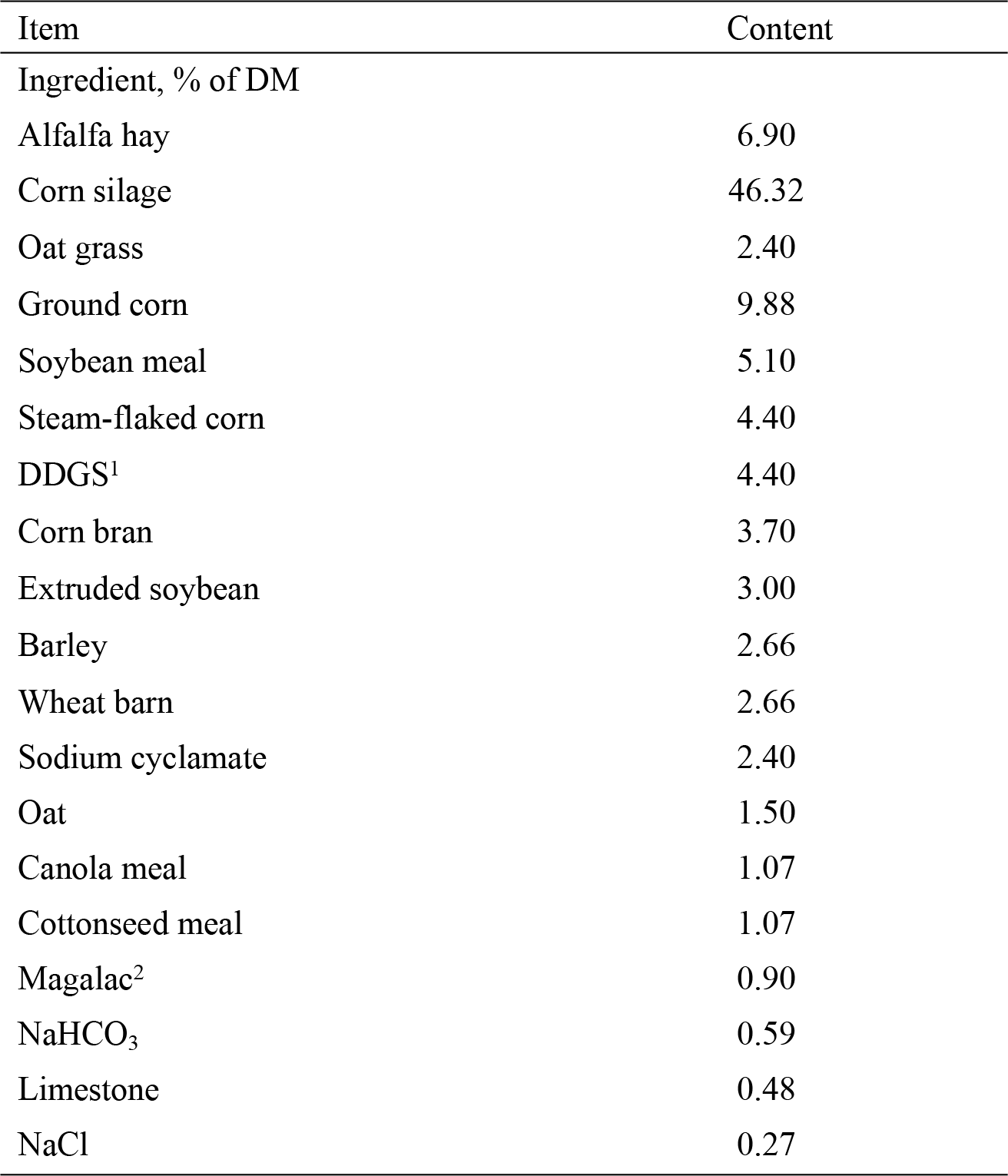
Ingredients and nutrient composition (% of DM) of the basal diet.

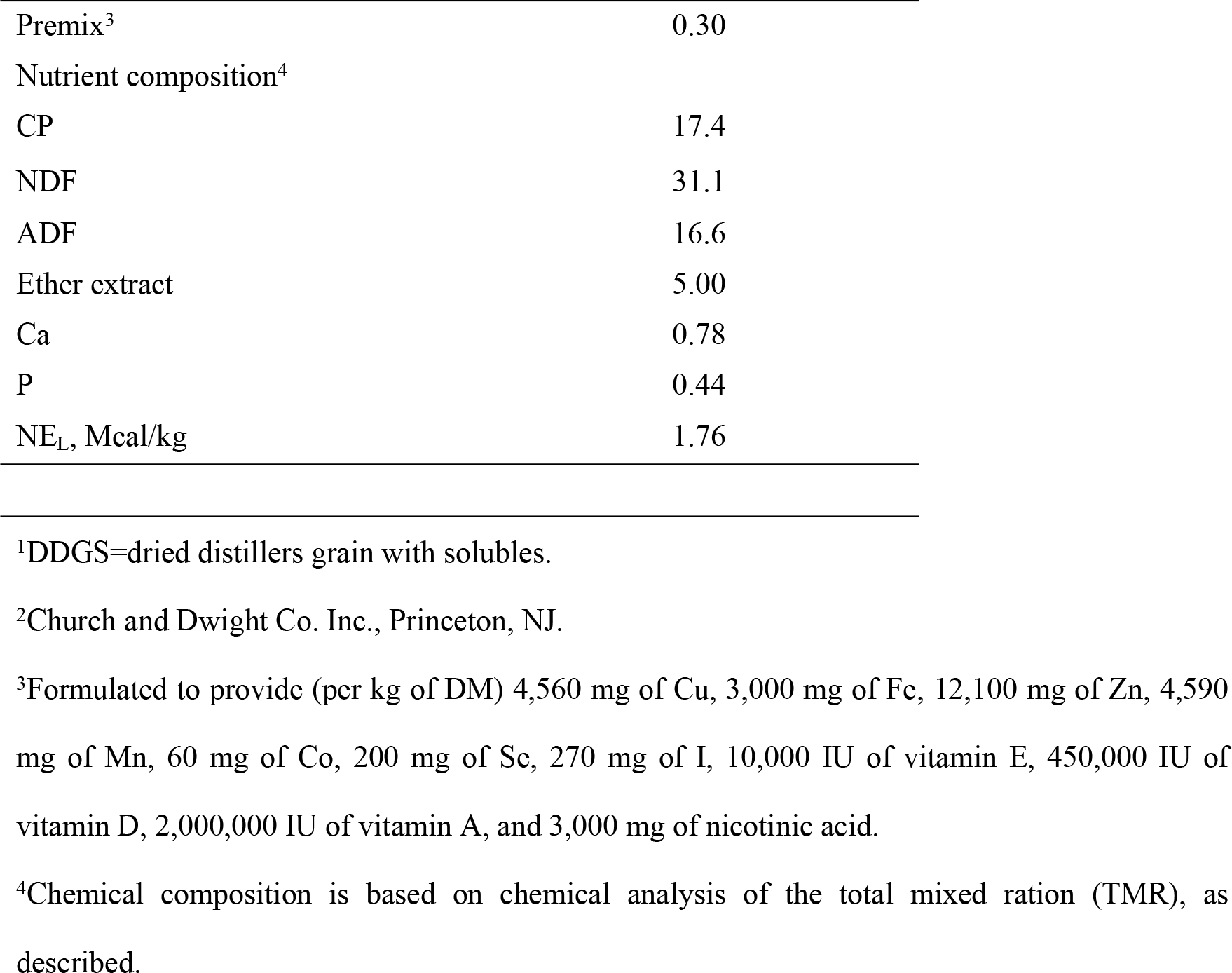

### Rumen fluid sampling and parameter measurement

Rumen fluid samples were collected from the oral cavity at 3-4 h after the morning feeding on day 7. The rumen contents were strained through 4 layers of cheesecloth with a mesh size of 250 μm. Ruminal pH was immediately measured using a portable pH meter (Testo 205, Testo AG, Germany). The filtered rumen fluid samples were centrifuged at 10,000 × g for 15 min at 4°C, aliquoted into 5-mL cryopreservation tubes, frozen in liquid nitrogen tank and stored at −80°C for analysis of the ruminal bacterial community. Another 10 mL of clear supernatant samples was mixed with 2 mL of 250 g/L of metaphosphoric acid and stored at −20°C for VFA determination, as described by Pan et al. [23].

### DNA extraction and polymerase chain reaction (PCR) amplification

Microbial DNA was extracted from rumen fluid samples using E.Z.N.A.® Bacterial DNA Kit (Omega Bio-Tek, Norcross, U.S.) according to the manufacturer’s protocols. The yield and purity of the extracted DNA were assessed with a NanoDrop 1000 instrument (NanoDrop, Wilmington, DE).

### 16S rRNA analysis

The V3-V4 regions of the 16S ribosomal RNA gene were amplified by PCR (95°C for 3 min, followed by 27 cycles at 95°C for 30 s, 55°C for 30 s, and 72°C for 45 s and a final extension at 72°C for 10 min) using the primers 338F 5’-barcode-ACTCCTACGGGAGGCAGCAG)-3’ and 806R 5’-GGACTACHVGGGTWTCTAAT-3’, where the barcode is an eight-base sequence unique to each sample. The PCR reactions were performed in triplicate in a 20-μL mixture containing 4 μL of 5 × FastPfu Buffer, 2 μL of 2.5 mM deoxyribonucleotide triphosphates (dNTPs), 0.8 μL of each primer (5 μM), 0.4 μL of FastPfu Polymerase, 0.2 μL of bovine serum albumin (BSA) and 10 ng of the template DNA. Amplicons were excised from 2% agarose gels and purified using AxyPrep DNA Gel Extraction Kit (Axygen Biosciences, Union City, U.S.) according to the manufacturer’s instructions and then quantified using QuantiFluor™ -ST (Promega, U.S.). Purified amplicons were pooled in equimolar ratios and pair-end sequenced (2 × 300) on the Illumina MiSeq platform according to standard protocols. The raw reads were deposited into the NCBI Sequence Read Archive (SRA) database (Accession Number: SRP136923).

The raw fastQ files were quality filtered by Trimmomatic and merged by FLASH under the following criteria. First, the reads were truncated at any site receiving an average quality score of <20 over a 50-bp sliding window. Second, sequences with overlapping segments longer than 10 bp were merged according to their overlapping region, with a mismatch of no more than 2 bp. Finally, sequences of each sample were separated according to barcodes (exactly matching) and primers (allowing 2 nucleotide mismatching), and reads containing ambiguous bases were removed.

Operational taxonomic units (OTUs) were clustered with a 97% similarity cutoff using UPARSE (version 7.1 http://drive5.com/uparse/) with a novel ‘greedy’ algorithm that simultaneously performs chimaera filtering and OTU clustering. The taxonomy of each 16S rRNA gene sequence was analysed according to the RDP Classifier algorithm (http://rdp.cme.msu.edu/) against the Silva (SSU123) 16S rRNA database using a confidence threshold of 70%.

### Statistical analysis

Data were analysed with a mixed-model procedure (SAS Institute, Inc., Cary, NC), and Duncan’s multiple comparison tests were employed to assess differences between means. Differences were considered statistically significant when *P*<0.05 and considered a trend when *P*<0.10.

## Results

### Dry matter intake, milk yield, and milk composition

Dry matter intake (DMI) was significantly greater in the high-yield (HY) group than in the low-yield (LY) group (*P*=0.03). Milk production (19.3±1.76 and 31.9±.76 kg/d, respectively; *P*<0.01), 4% fat-corrected milk (FCM) (18.95±1.73 and 29.20±1.73 kg/d, respectively. *P*<0.01) and energy-corrected milk (ECM) (21.07±1.93 and 32.09±1.93 kg/d, respectively; *P*<0.01) were significantly lower in the LY group than in the HY group (Table 2). The milk fat content tended to be higher (*P*=0.08) (Table 2), and the milk protein content was significantly higher in the LY group than in the HY group (*P*<0.01). No difference was observed in milk lactose content between the LY and HY groups (*P*=0.21; Table 2). Fat, protein and lactose yields were significantly greater in the HY group than in the LY group (Table 2). However, somatic cell count (SCC) was not different between the HY and LY groups (*P*=0.13; Table 2).

**Table 2.**
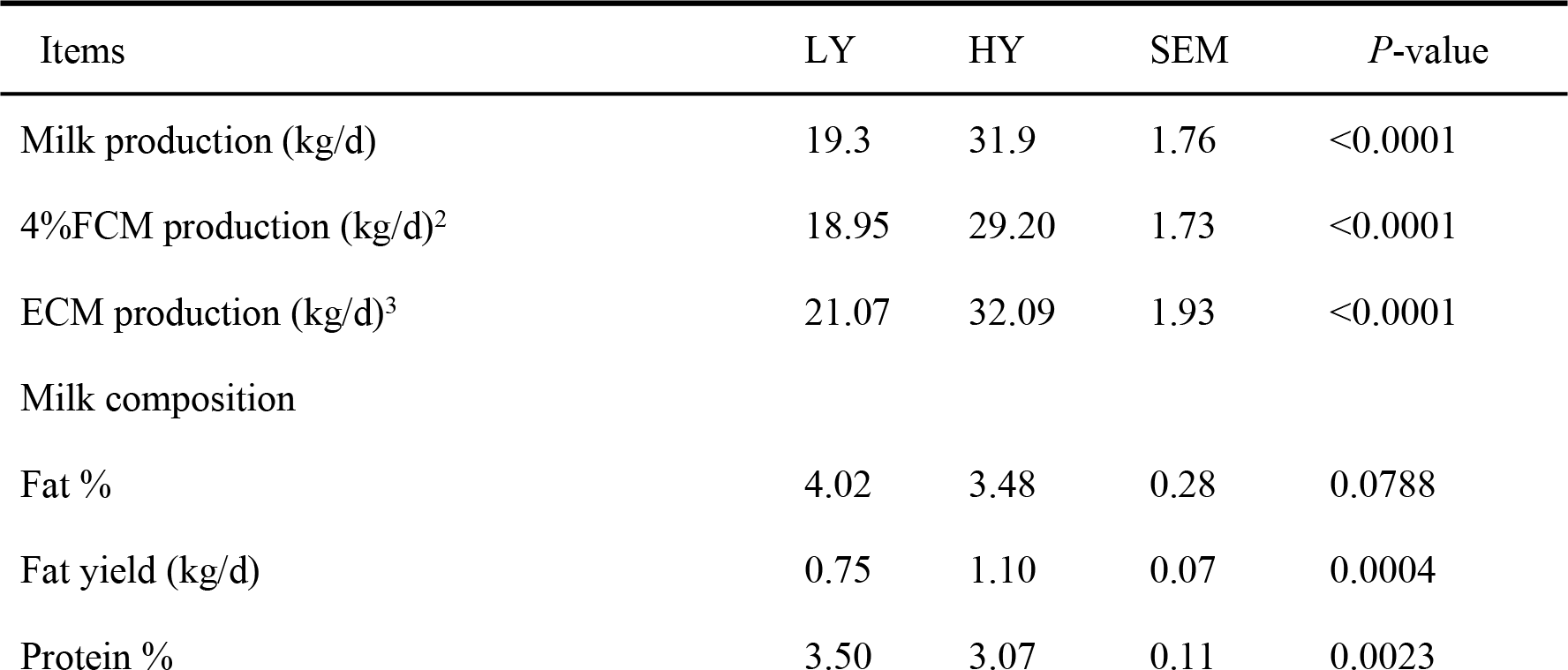
Milk and ECM from high-yielding and low-yielding dairy cows during the entire sampling period ^1^.

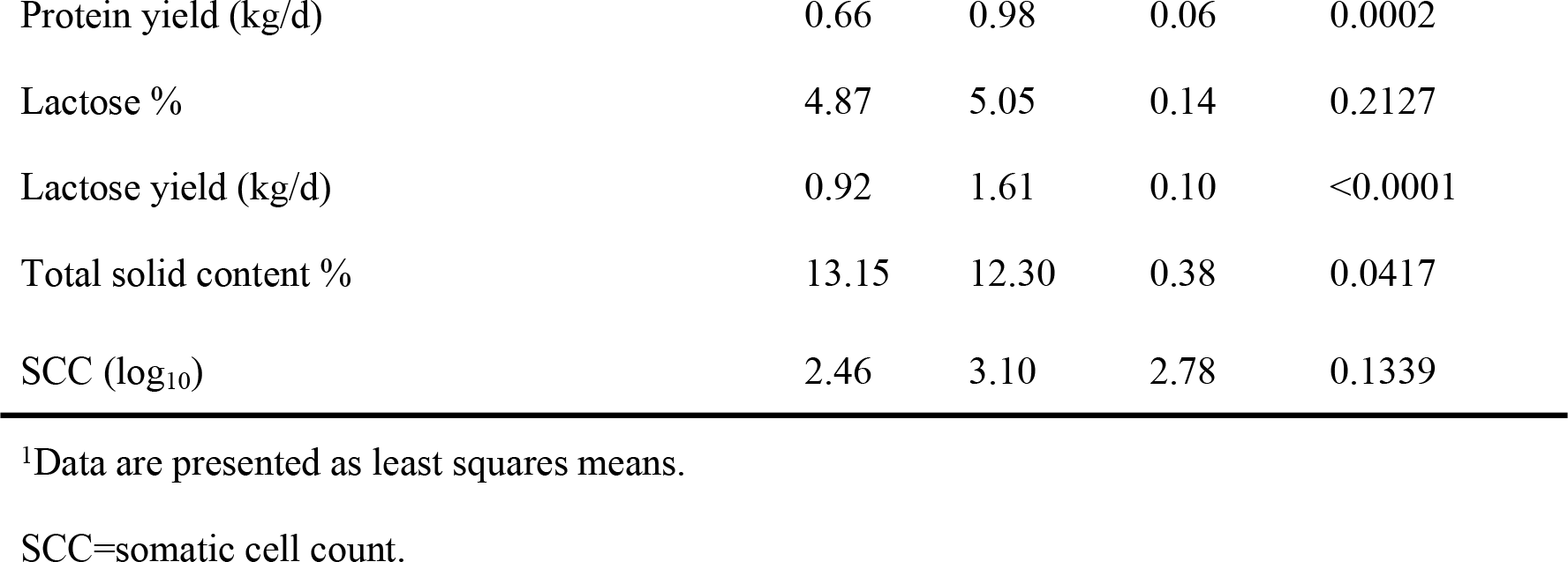

### Ruminal pH and VFA concentrations

Rumen pH was not different between the HY group and the LY group (Table 3). NH_3_-N (mg/dL) was greater in the HY group than in the LY group (*P*<0.01), and there was a significant increase in acetate, propionate and butyrate levels in the HY group relative to the LY group (*P*<0.05).

In addition, total VFA levels were significantly greater in the HY group than in the LY group (*P*=0.02). A lower trend was observed for the acetate:propionate ratio in the HY group compared to the LY group (*P*=0.06).

**Table 3.** Effects of differences between high-yielding and low-yielding dairy cows on metabolites in the rumen.

| Items | LY | HY | SEM | *P*-value |
| --- | --- | --- | --- | --- |
| pH | 6.73 | 6.71 | 0.02 | 0.69 |
| NH<sub>3</sub>-N, mg/dL | 7.99 | 13.28** | 1.03 | 0.01 |
| **Molar proportion** |  |  |  |  |
| Acetate | 62.19 | 68.97* | 1.78 | 0.05 |
| Propionate | 21.94 | 26.35** | 0.97 | 0.02 |
| Isobutyrate | 0.83 | 0.84 | 0.03 | 0.93 |
| Butyrate | 12.01 | 14.23** | 0.45 | 0.01 |
| Isovalerate | 1.22 | 1.48 | 0.07 | 0.08 |
| Valerate | 1.57 | 1.76 | 0.06 | 0.14 |
| Acetate:propionate ratio | 2.85 | 2.64 | 0.06 | 0.06 |
| Total VFA, mM | 99.76 | 113.63 | 3.14 | 0.02 |

\* $P<0.05$ ; \*\* $P<0.01$ : Values within a sampling day followed by superscripted asterisks differ.
SEM=standard error of the mean.

### Diversity and richness of microbial communities

In total, 2,382,338 merged sequences were acquired for the 16 samples from the dairy cows, and 1,191,169 high-quality sequences, with an average read length of 440 bp, were classified as bacterial. On average, at least 54,144 sequences were obtained per sample, and greater than 99% depth coverage was achieved. The rarefaction curve generated tended to plateau, showing that the number of OTUs did not rise with an increasing volume of data. This finding showed that the data volume of sequencing was reasonable. The results of this study show that the sequencing data were reasonable and could reflect changes in most bacterial flora (Figure 1).

**Figure 1.**
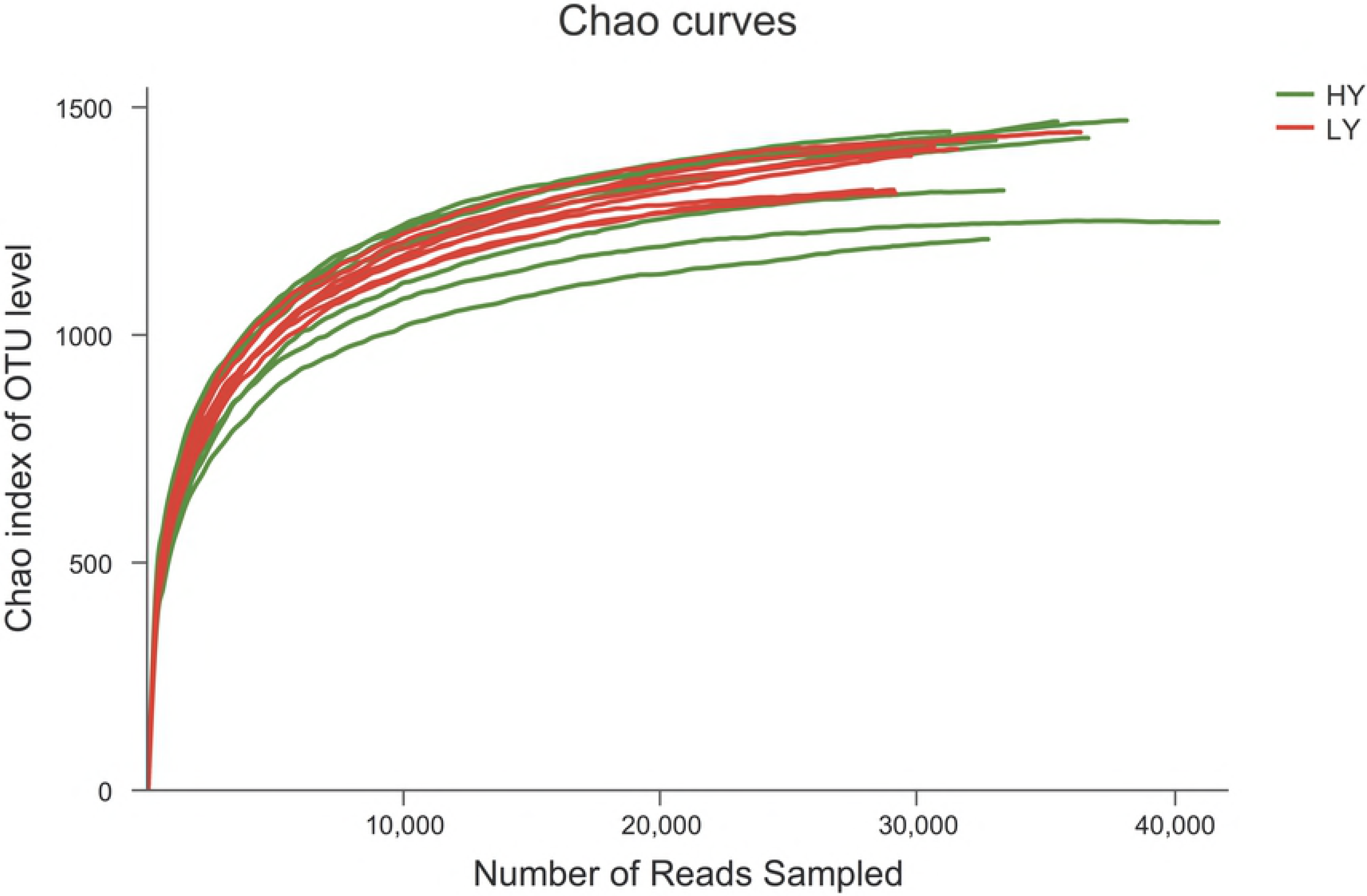
The rarefaction curve of sequencing based on OTUs^ [1 OTUs, operational taxonomic units.]^.

No significant differences were observed in alpha diversity index results between the HY and LY groups (*P*>0.05) (Table 4). However, the coverage of the HY group was significantly higher than that of the LY group (*P*<0.01), indicating greater community diversity for the HY group.

**Table 4.** Alpha diversity index of rumen bacteria.

| Item | LY | HY | SEM | *P*-value |
| --- | --- | --- | --- | --- |
| Sobs | 1196.13 | 1214.88 | 17.21 | 0.60 |
| Shannon | 5.4104 | 5.4852 | 0.04 | 0.42 |
| Simpson | 0.01 | 0.01 | 0.00 | 0.74 |
| ACE | 1375.65 | 1360.65 | 18.75 | 0.70 |
| Chao | 1378.64 | 1376.82 | 20.10 | 0.97 |
| Coverage | 0.9921 | 0.9938** | 0.00 | 0.001 |

\* $P<0.05$ ; \*\* $P<0.01$ : Values within a sampling day followed by superscript asterisks differ.
SEM=standard error of the mean;

Principal coordinate analysis (PCoA) results showed that the HY group was distinct from the LY group (Figure 2). Principal coordinates 1 and 2 accounted for 24.94% and 12.25%, respectively, of the total variation.

**Figure 2.**
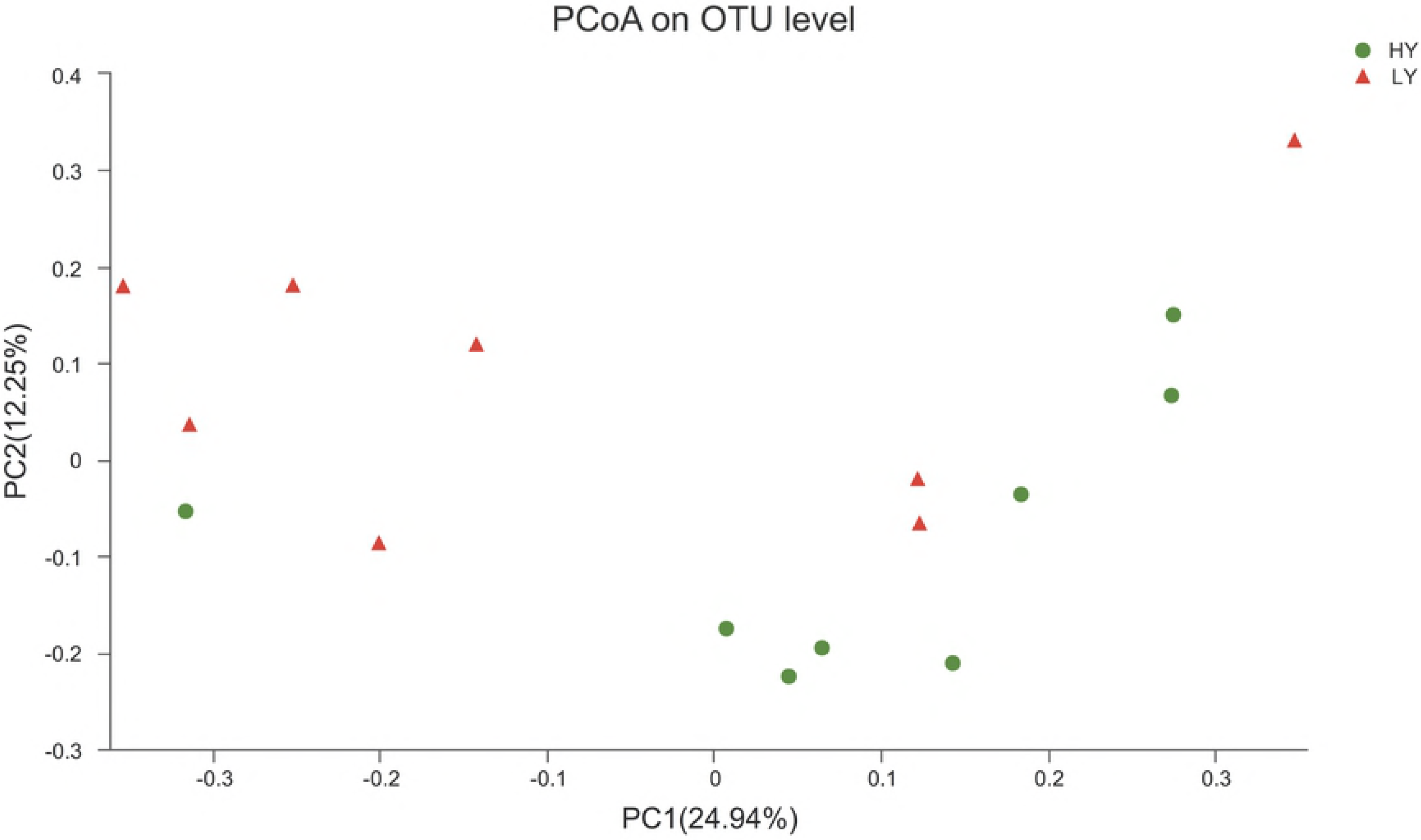
Principal coordinate analysis (PCoA) of bacterial community structures of the ruminal microbiota in the high-yielding group (green circles) and the low-yielding group (red triangles).PCoA plots were constructed using the unweighted UniFrac method.

Twenty-one bacterial phyla were identified across all samples. Bacteroidetes, Firmicutes and Proteobacteria were the three dominant phyla, representing 57.59%, 35.86%, and 1.53%, respectively, of the total sequences (Figure 3). Thus, at the phylum level, Bacteroidetes and Firmicutes were particularly dominant. The HY group exhibited a greater abundance of Firmicutes and lower abundance of Bacteroidetes than did the LY group (*P*<0.01), whereas Proteobacteria was less abundant (*P*<0.05).

**Figure 3.**
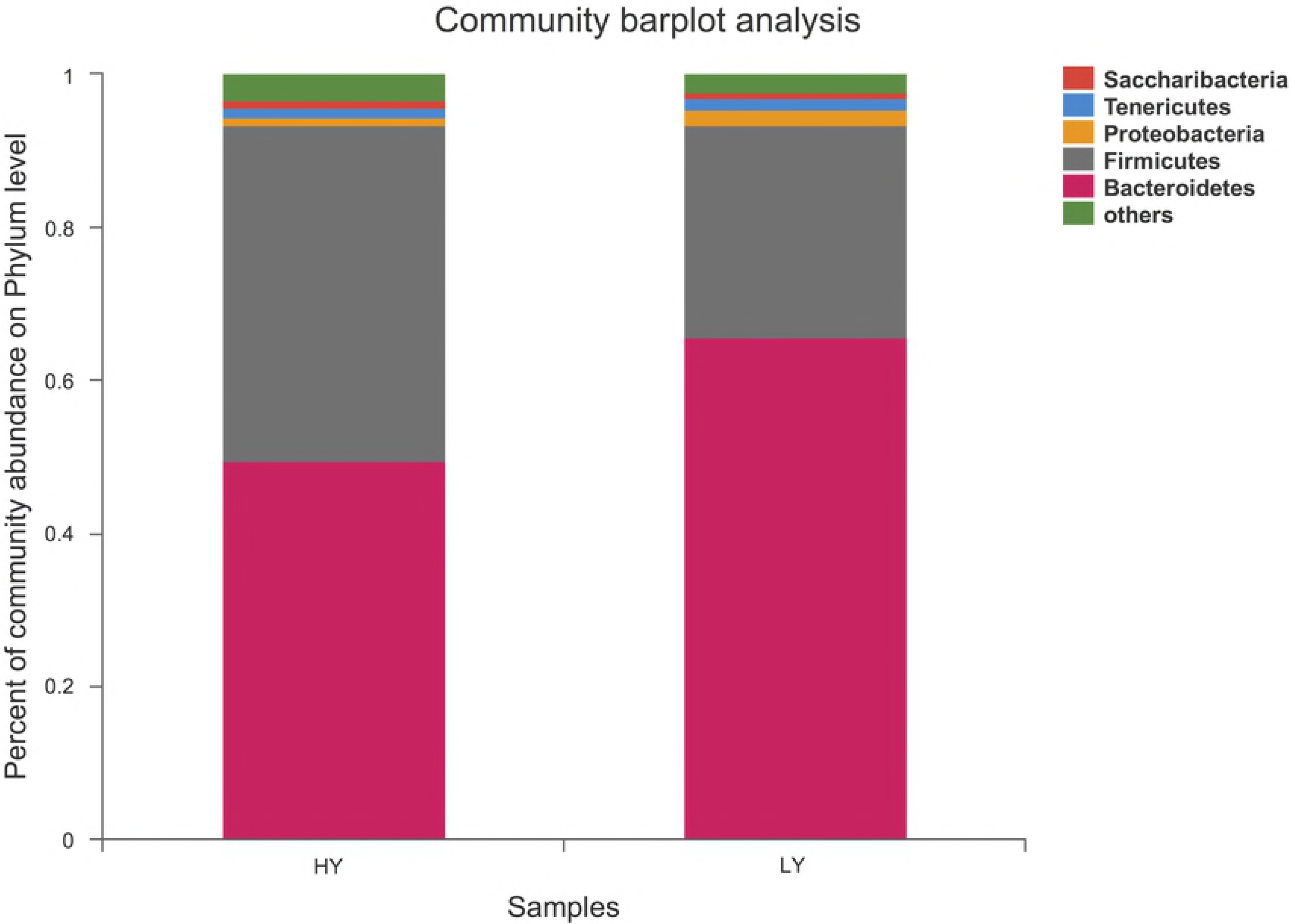
Percent composition of predominant phyla in the rumen.

At the genus level, taxa with a relative abundance of ≥1% in at least one sample were further analysed, and the relevant genera are presented in Figure 4 and Figure 5. Twenty-one genera were identified, 6 of which exhibited significantly different abundances between the groups. Specifically, 4 genera were more abundant in the HY group at *P*<0.01, including *Ruminococcaceae-NK4A214-group, Ruminococcus 2*, *Lachnospiraceae-BS11-gut-group*, and *[Eubacterium]-coprostanoligenes-group*, and 2 were more abundant in the HY group at *P*<0.05: *Succiniclasticum* and *Christensenellaceae-R-7-group*.

**Figure 4.**
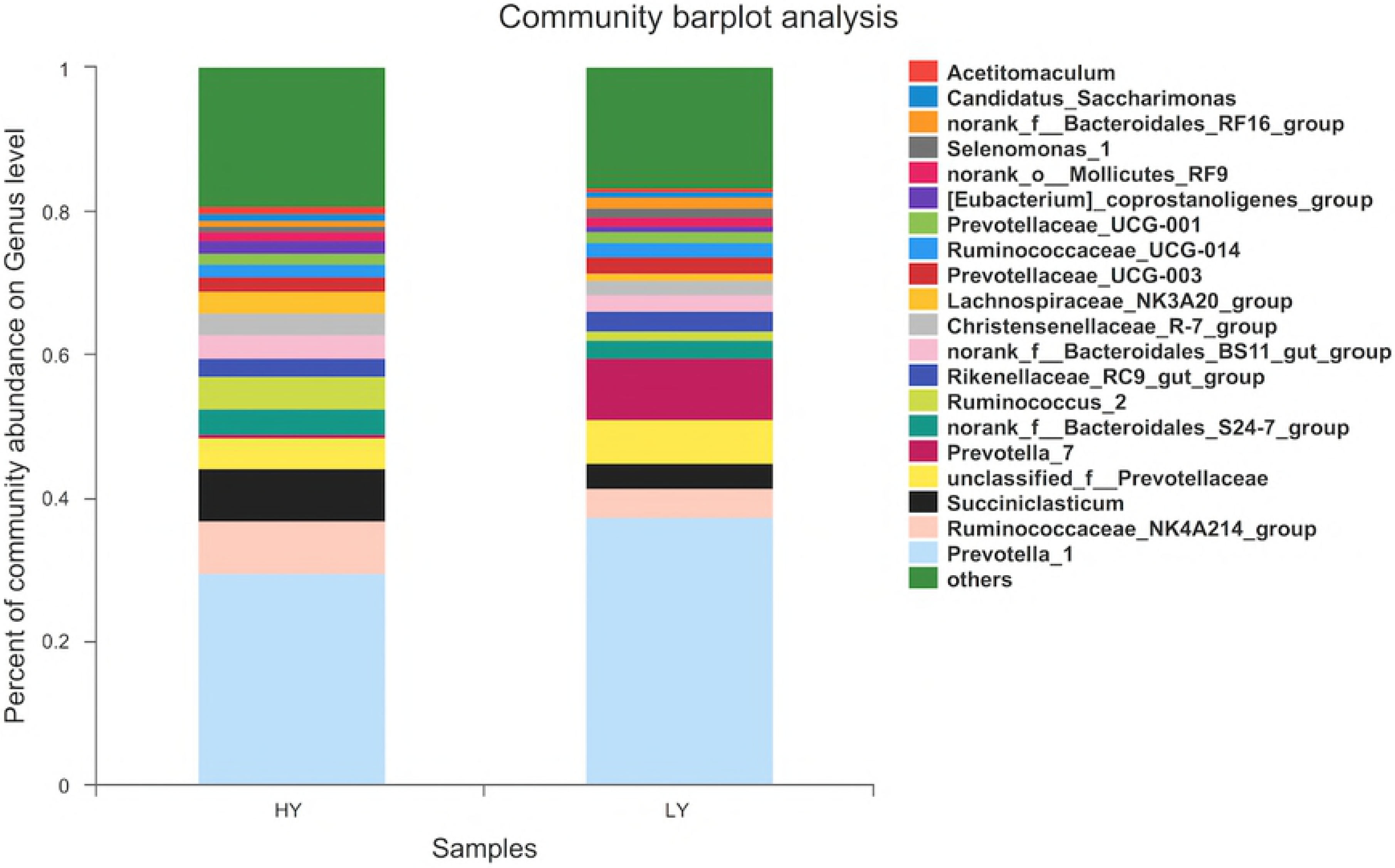
Percent composition of genera in the rumen.

**Figure 5.**
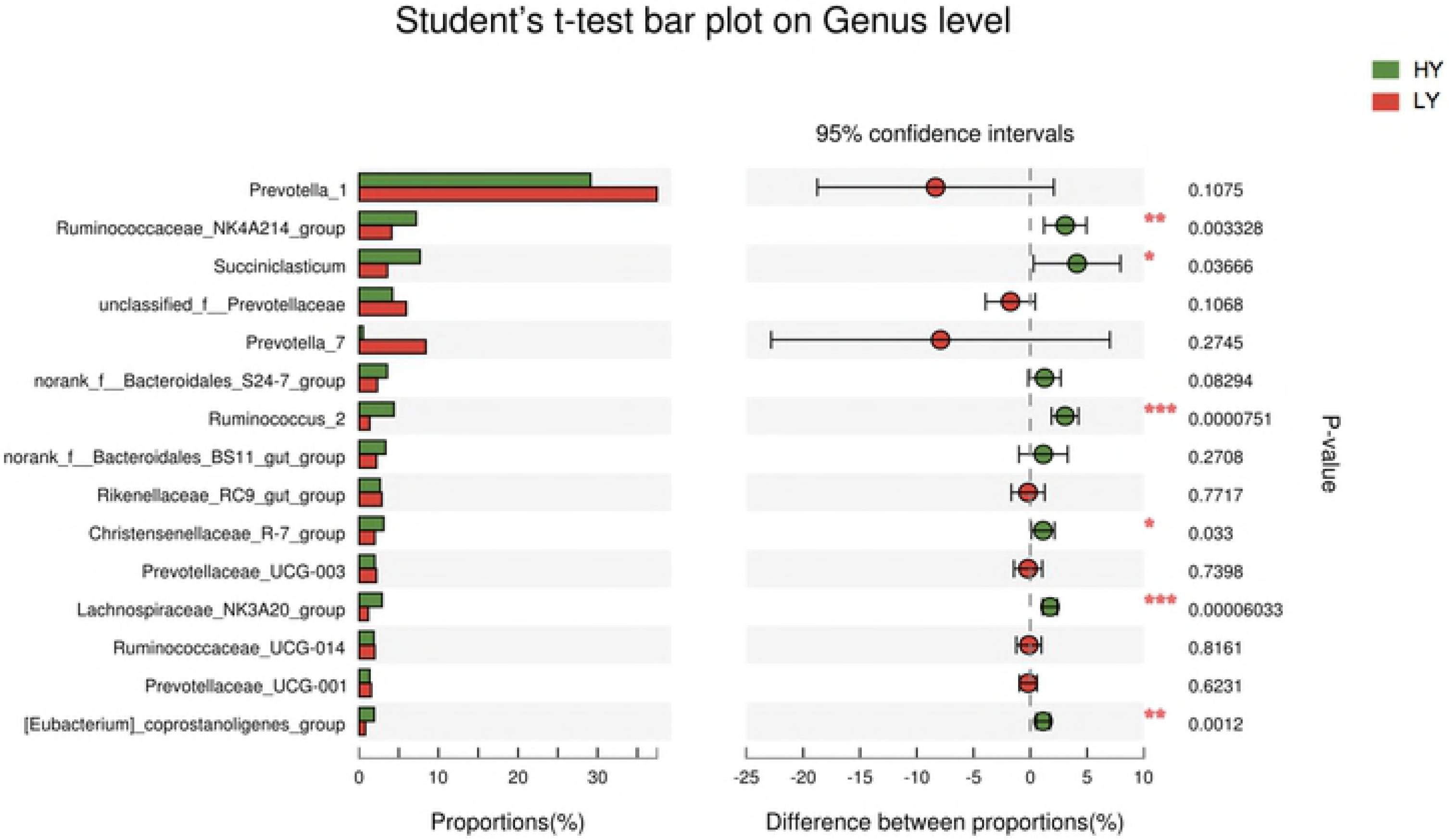
Percent composition and significance of genera in the rumen.

### Correlations between bacterial communities and ruminal variables

As shown in Figure 6, the relative abundances of the genera *Bacteroides* and *Ruminococcus 2* were positively correlated with ruminal propionate and NH_3_-N concentrations (r>0.4, *P*<0.05) but negatively correlated with the ruminal ratio (acetate:propionate ratio) (r<−0.4, *P*<0.05). In addition, *norank_o_Mollicutes_RF9* was positively correlated with ruminal acetate and VFA concentrations (r>0.4, *P*<0.05). *Candidatus-Saccharimonas* was positively correlated with the ruminal propionate concentration (r>0.4, *P*<0.05) but negatively correlated with the ruminal ratio (r<−0.4, *P*<0.05). Moreover, the ratio was negatively correlated with *Schwartzia* (r<−0.6, *P*<0.05).

**Figure 6.**
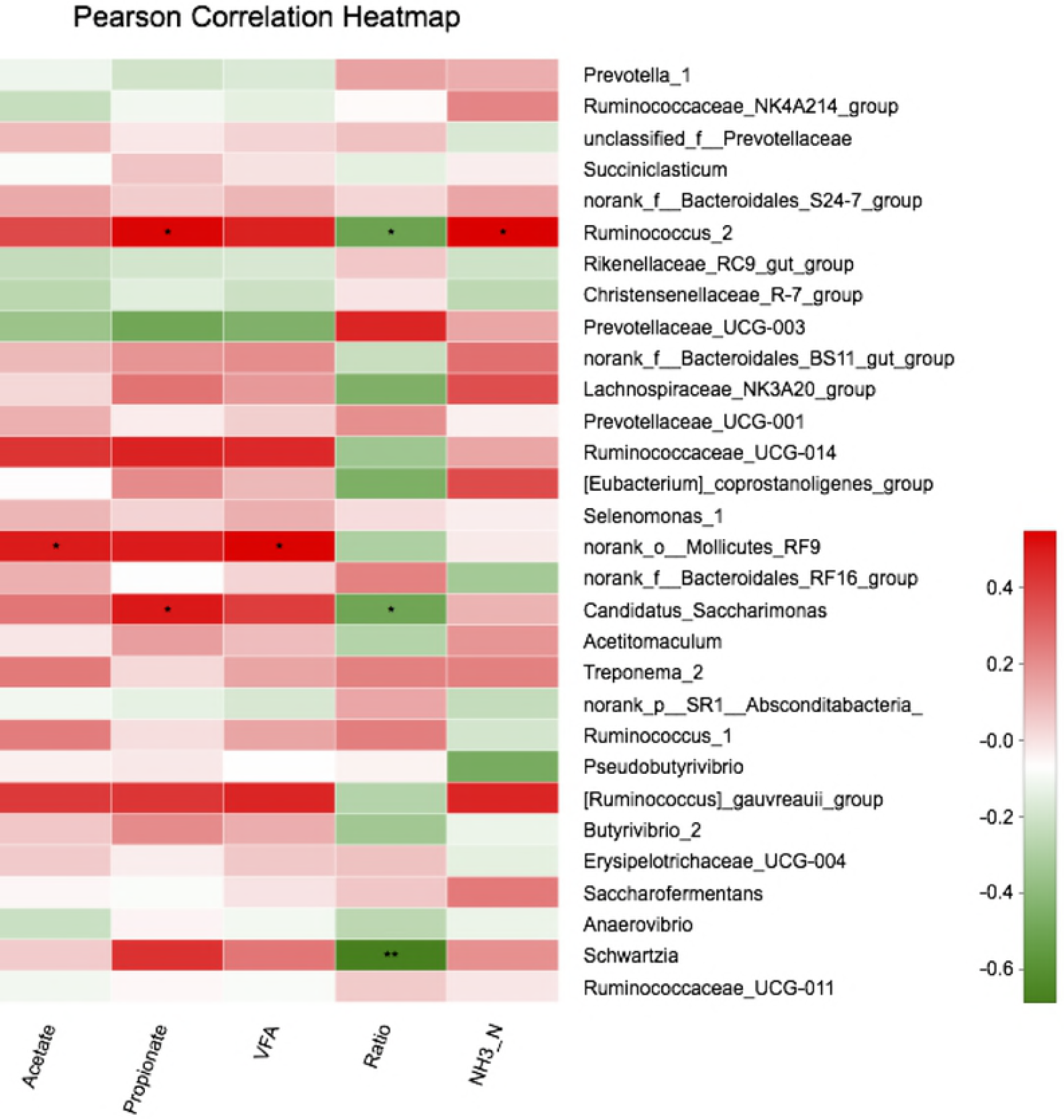
Correlation analyses between the relative abundances of bacteria genera and ruminal fermentation parameters. Only genera with abundances significantly associated with ruminal VFA, propionate and acetate concentrations are presented. Green represents a negative correlation between the abundance of the species and the ratio (r<−0.4), and red represents a positive correlation (r>0.4, 0.01<*P*<=0.05 *; 0.001<*P*≤0.01 **; P≤0.001 *** Values with a significant correlation followed by superscripted asterisks differ).

## Discussion

High-throughput sequencing has been widely used in studies of microbial flora in ruminants as a means of quickly and efficiently determining the microbial community structure of the rumen [24]. Differences in rumen microbiota between high-and low-yield dairy cows can therefore be characterised using high-throughput sequencing methods. The present study also linked rumen fluid VFAs with milk components and rumen microbes.

### Changes in rumen fermentation parameters and milk composition

Higher values for rumen fermentation parameters and milk composition were found in the HY group compared to the LY group. This finding suggests that due to host-microbiota interactions, different cows may harbour different microbial species compositions, which are closely related to distinct differences in rumen fermentation parameters and milk composition. Furthermore, recent findings suggest an important relationship between VFAs and milk components [2, 25, 26]. Specifically, of the three principle VFAs, acetic and butyric acids are substrates for oxidation and are precursors of lipids [27, 28]. Propionic acid is the only glucogenic VFA, accounting for 65-80% of the net glucose supply in lactating dairy cows [29, 30]. In a previous study, milk yield was most highly related to rumen concentrations of butyrate and propionate [31]. In the present study, propionate and butyrate concentrations as well as fat, protein and lactose yields were significantly greater in the HY group than in the LY group, which is consistent with previous research [32]. The NH_3_-N concentration in rumen fluid can reflect the balance of protein degradation and synthesis under varying feed conditions. NH_3_-N is an intermediate product of feed protein, nonprotein nitrogen degradation and microbial protein synthesis, and it is mainly affected by feed protein degradation, rumen wall absorption, microorganism utilisation and rumen chyme outflow rate [33–35]. Yang et al. [36] reported that the concentration of NH_3_-N should be higher than 5 mg/dL; otherwise, it will influence the “uncoupling” effect of ruminal fermentation and reduce the efficiency of microbial protein synthesis. According to our results, the NH_3_-N concentration was within the normal range, though that in the HY group was significantly higher than that in the LY group. These results indicate that rumen microbes promote protein degradation, providing a better understanding of difference in milk proteins between the two groups.

### Differences in rumen microbial composition between HY and LY groups

No differences were observed in bacterial community richness and diversity between the groups. Three phyla predominated in both groups, which was consistent with previous studies reporting that the principal phyla of microbes in the rumen are Bacteroidetes, Firmicutes, and Proteobacteria. The proportions of these three phyla account for approximately 94% of the total [35, 37, 38]. Interestingly, our results showed that the abundance of Firmicutes in the HY group was higher than that in the LY group, though the abundances of the two other dominant phyla were lower in the former than in the latter.

As previously described by Pan et al. [23], cows fed a high proportion of grain have a higher abundance of Firmicutes and a lower abundance of Proteobacteria than control cows, and other studies have shown that feeding a high amount of grain can promote milk production [39, 40]. Thus, the present study provides a better understanding of why cows fed the same diet can have different milk production. The present findings further demonstrate that Firmicutes plays an important role in milk production.

In agreement with other research results [11, 41], *Prevotella* was the most abundant genus in all samples. Although *Prevotella* was more abundant in the LY group than in the HY group, the difference was not significant. In contrast, *Ruminococcaceae-NK4A214-group, Ruminococcus 2*, *Lachnospiraceae-BS11-gut-group* and *[Eubacterium]-coprostanoligenes-group* were significantly different between the two groups, with higher abundances in the HY group than in the LY group. Jiang et al. [42] reported that the increase in the relative abundance of *Ruminococcus* partly explains why adding live yeast to the diet increases the in vivo digestibility of DM and NDF and the performance of cows. This result illustrates that high-performance cows have higher abundances of *Ruminococcus* in the rumen fluid, which is consistent with the present research results. Members of the family Lachnospiraceae are gram-positive obligate anaerobes that are mostly non-spore-forming bacteria [43, 44]. Huws et al. [45] showed that Ruminococcaceae and Lachnospiraceae play predominant roles in biohydrogenation pathways within the rumen. As the primary succinate-utilising bacterial taxon, *Succiniclasticum* accounted for 7.45% of the total bacterial community in the HY group, with significantly greater abundance than in the LY group. A higher level of *Succiniclasticum* has been associated with greater production of succinate from starch degradation [46]. Moreover, the abundances of Christensenellaceae and Ruminococcaceae NK4A214 in the HY group were significantly higher than in the LY group, though little information about these two genera has been reported in the literature. The reasons for the altered status of genera in cows with different milk production are unclear.

## Conclusion

In summary, high-yield dairy cows have better ruminal fermentation patterns than do low-yield cows, which was partially attributed to the greater abundances of *Bacteroides, Ruminococcus 2, Ruminococcaceae NK4A214*, Lachnospiraceae, *Succiniclasticum, Eubacterium* and *Christensenella* in the former. Furthermore, rumen fermentation in high-yield cows exhibited higher VFA levels than that found in low-yield cows. Rumen microbiotal composition between high-yield and low-yield dairy cows differs, and microbial species diversity and distribution contribute to production-related phenotypes. Overall, our findings enhance our understanding of rumen bacteria in cows with different milk yields and provide new strategies for improving dairy cow production performance.

## Acknowledgements

The study was supported by the National Nature Science Foundation of China (Grant No. 31772629 and No. 31702302), Beijing Municipal Education Commission Project (SQKM201710020011), Open Project Program of Beijing Key Laboratory of Dairy Cow Nutrition, Beijing University of Agriculture and the National Key Research and Development Plan (2016YFD0700205, 2016YFD0700201, 2017YFD0701604). The authors thank the members of the Beijing Key Laboratory for Dairy Cow Nutrition, Beijing University of Agriculture (Beijing, China) for their assistance in the sampling and analysis of the samples.

